# PDBe tools for an in-depth analysis of small molecules in the Protein Data Bank

**DOI:** 10.1101/2024.11.05.622044

**Authors:** Preeti Choudhary, Ibrahim Roshan Kunnakkattu, Sreenath Nair, Dare Kayode Lawal, Ivanna Pidruchna, Marcelo Querino Lima Afonso, Jennifer R. Fleming, Sameer Velankar

## Abstract

The Protein Data Bank (PDB) is the primary global repository for experimentally determined 3D structures of biological macromolecules and their complexes with ligands, proteins, and nucleic acids. PDB contains over 47,000 unique small molecules bound to the macromolecules. Despite the extensive data available, the complexity of small molecule data in the PDB necessitates specialised tools for effective analysis and visualisation. PDBe has developed a number of tools, including PDBe CCDUtils (https://github.com/PDBeurope/ccdutils) for accessing and enriching ligand data, PDBe Arpeggio (https://github.com/PDBeurope/arpeggio) for analysing interactions between ligands and macromolecules, and PDBe RelLig (https://github.com/PDBeurope/rellig) for identifying the functional roles of ligands (such as reactants, cofactors, or drug-like molecules) within protein-ligand complexes. Furthermore, the enhanced ligand annotations and data generated by these tools are presented in a comprehensive view on the novel PDBe-KB ligand pages, providing a holistic view of small molecules that enables the establishment of their biological contexts (Example page for Imatinib: https://wwwdev.ebi.ac.uk/pdbe-srv/pdbechem/chemicalCompound/show/STI). By improving the standardisation of ligand identification, adding various annotations, and offering advanced visualisation capabilities, these tools help researchers navigate the complexities of small molecules and their roles in biological systems, facilitating mechanistic understanding of biological functions. The ongoing enhancements to these resources are designed to support the scientific community in gaining valuable insights into ligands and their applications across various fields, including drug discovery, molecular biology, systems biology, structural biology, and pharmacology.

## Introduction

Protein Data Bank in Europe (PDBe) is one of the founding partners of the worldwide Protein Data Bank (wwPDB) consortium, dedicated to collecting, curating, and providing access to a global repository of macromolecular structure models, the Protein Data Bank (PDB) (wwPDB consortium 2019). The PDB contains over 225,000 experimentally determined macromolecular structures, with around 75% featuring at least one bound small molecule. These small molecules serve various purposes, including experimental necessities or acting as ligands with diverse biological functions, like cofactors, metabolites, substrates, and inhibitors. To ensure a standardised representation of small molecules, the wwPDB developed the Chemical Component Dictionary (CCD), a comprehensive reference resource that includes data for all unique chemical components found within PDB entries, including individual amino acids, nucleotides, and ligands (Westbrook et al. 2015). The CCD provides details such as chemical descriptors (e.g., chemical formula, molecular weight, SMILES, and InChI), systematic chemical names, chemical connectivities, stereochemical assignments, and idealised 3D coordinates generated using Molecular Networks’ Corina (Schwab 2010) and OpenEye’s OMEGA (Hawkins et al. 2010). The CCD is accessible at https://www.wwpdb.org/data/ccd. Each unique chemical component is assigned a CCD identifier, allowing for the identification of all instances of a specific small molecule in PDB structures. For instance, adenosine triphosphate has a CCD identifier: ATP, and is bound in over 3,600 macromolecular structures.

Complex ligands are often fragmented into individual chemical components during the refinement and biocuration, creating challenges for their identification and mapping to other databases. To address this, the wwPDB introduced the Biologically Interesting Molecule Reference Dictionary (BIRD) in 2013 (Dutta et al. 2014). This manually curated reference dictionary assigns unique identifiers and detailed descriptions to peptide-like inhibitors, antibiotic molecules and common oligosaccharides found in PDB entries. Each entry details the composition, connectivity, chemical structure, and functions of the reference molecule. The BIRD reference data is accessible at https://www.wwpdb.org/data/bird, with examples like Vancomycin, identified by the BIRD identifier: PRD_000204. In 2020, the wwPDB standardised the representation of carbohydrate polymers, which had been previously fragmented into individual monosaccharides, by introducing a new ’branched’ entity representation (Shao et al. 2021). This improved system offers a more accurate depiction of complex carbohydrate structures and is supported by consistent 2D representations based on the Symbol Nomenclature for Glycans (SNFG) (Neelamegham et al. 2019). While these wwPDB efforts have significantly improved the handling of peptide-like inhibitors, antibiotics, and carbohydrates, several multi-component ligands remain fragmented into separate components.

To comprehensively tackle complex ligands fragmentation, PDBe introduced Covalently Linked Components (CLCs) (Kunnakkattu et al. 2023). This novel class of reference small molecules identifies ligands composed of multiple covalently linked Chemical Components (CCDs) across the entire PDB archive. CLCs provide a more complete and accurate representation of these complex ligands, filling gaps left by fragmented CCDs not included in the BIRD or carbohydrate remediation efforts. To streamline identification and analysis of CLCs, PDBe has implemented an automatic process to assign unique identifiers based on InChIKey. For instance, in PDB entry 1D83, two carbohydrates. Carbohydrate 1: 2,6-dideoxy-4-O-methyl-alpha-D-galactopyranose-(1-3)-(2R,3R,6R)-6-hydroxy-2-methyltetrahydro-2H-pyran-3-yl acetate, composed of subcomponent CCDs: CDR, CDR, and ERI. Carbohydrate 2: 3-C-methyl-4-O-acetyl-alpha-L-olivopyranose-(1-3)-(2R,5S,6R)-6-methyltetrahydro-2 H-pyran-2,5-diol-(1-3)-(2R,5S,6R)-6-methyltetrahydro-2H-pyran-2,5-diol, consisting of subcomponent CCDs: ARI and 1GL. These two carbohydrates are covalently linked to the CCD: CPH to form a single small molecule, Chromomycin. This complex ligand, consisting of the covalently linked CCDs 1GL, ARI, CPH, CDR, CDR, and ERI, can now be identified as a single entity using the CLC identifier: CLC_000153 (Figure 1).

**Figure 1.**
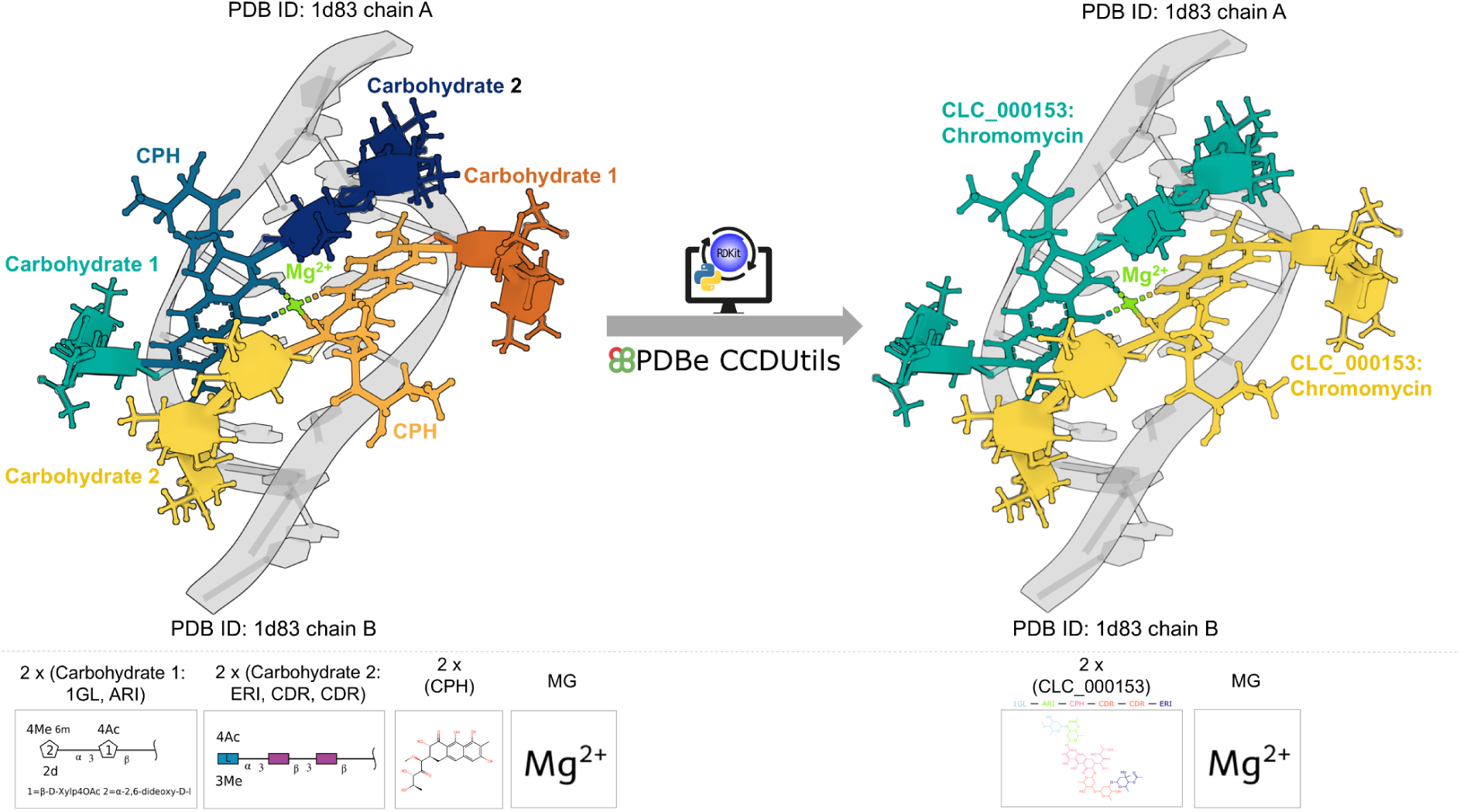
Structure of Chromomycin dimer/DNA oligomer complex (PDB ID 1D83). Originally annotated with four separate ligands per chain (two copies each of Carbohydrate 1, Carbohydrate 2, CPH, and Mg), the first three components are now unified as a single Chromomycin ligand (CLC_000153) per chain. The structure features a Chromomycin dimer coordinating with Mg.

Notably, BIRD entries are manually curated, and some CLCs may eventually transition into BIRD entries. For example, in PDB entry 1AO4, Peplomycin—a glycopeptide antineoplastic antibiotic—was previously fragmented into the CCDs PMY, GUP, and 3FM. However, it can now be represented as a single entity with the CLC identifier CLC_000034. If Peplomycin is later classified as a BIRD, CLC_000034 will be automatically replaced by the corresponding BIRD identifier. Creating CLCs to represent chemically complete ligands allows for improved mapping to other chemical databases such as PubChem, ChEMBL, and KEGG, overcoming the limitations of fragmented representation using multiple CCDs. Together, the CCD, BIRD (PRD) and CLC reference dictionaries provide a comprehensive set of unique small molecules found within the PDB, enhancing the understanding and exploration of small molecules bound to biological macromolecules.

The complexity and diversity of small molecules in the PDB necessitate the development of specialised tools for their access and analysis. Although existing small-molecule reference dictionaries provide essential data, additional tools are required to enhance this information within a broader biological context. Furthermore, it is necessary to elucidate the functional roles of ligands within these complexes and distinguish them from experimental artefacts, such as buffers and cryoprotectants. Accurately identifying complete ligand-macromolecule complexes and key protein-ligand interactions is essential for understanding protein functions. Here, we discuss PDBe’s strategies to address these challenges, explore PDBe tools and provide tutorials (Table 1) on their utilisation for ligand analysis and visualisation. These tools empower researchers to navigate the complexities of ligand and ligand-macromolecule complex structures, assess their biological significance, and perform comparative analyses across various databases (Figure 2).

**Figure 2.**
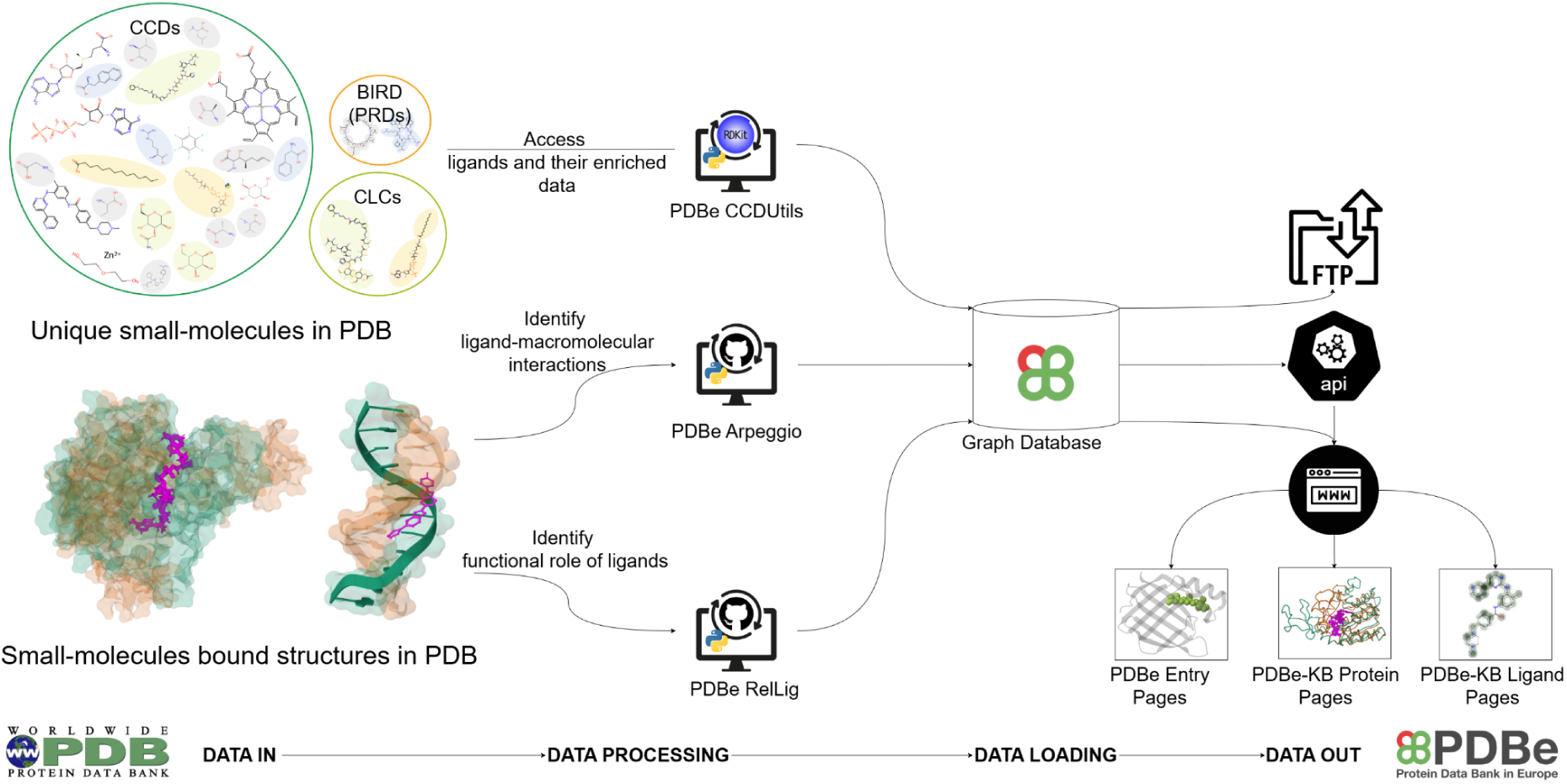
Schematic overview of the PDBe ligand tools for ligand analysis and visualisation. The PDBe CCDUtils allows access to small molecules, while PDBe Arpeggio identifies ligand-macromolecular interactions, and PDBe RelLig determines ligand functional roles in ligand-protein complexes. PDB data is processed weekly through these pipelines and loaded into the PDBe Graph database, supporting up-to-date data files on FTP, API endpoints, and various web pages, including PDBe entry pages and PDBe-KB Protein and Ligand pages.

**Table 1:**
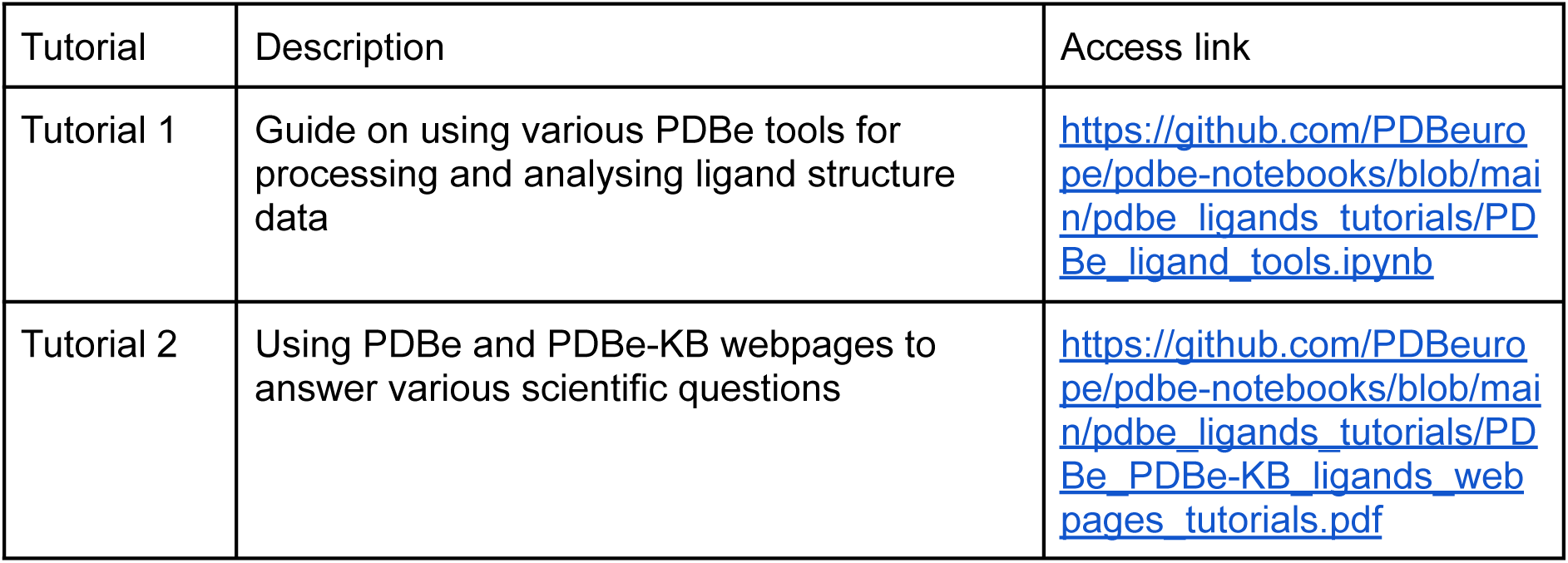
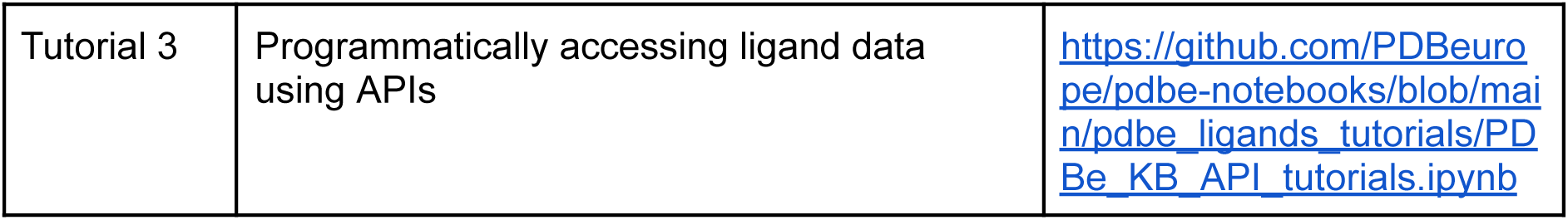
Tutorials demonstrating the use of PDBe tools for in-depth analysis and visualisation of ligands in PDB.

### PDBe tools for processing ligand structure data and their analysis

PDBe CCDUtils: Accessing ligand and their enriched data in PDB PDBe CCDUtils (Kunnakkattu et al. 2023) is an open-source chemistry toolkit designed to improve the parsing, processing, and analysis of small molecules within the PDB’s reference dictionaries, including CCD, PRD, and CLC. Built on RDKit (Landrum et al. 2024), it extends functionality by adding support for the PDBx/mmCIF file format (Westbrook et al. 2022) and provides access to variety of metadata such as chemical descriptors (SMILES, InChI, InChIKey), 2D depictions and 3D coordinates. It computes RDKit-derived physicochemical properties, generates 3D conformers and identifies core chemical substructures using scaffold detection methods like Murcko Scaffolds (Bemis and Murcko 1996) and BRICS (Degen et al. 2008). PDBe CCDUtils can also be used to search ligand substructures against a curated library of over 2,000 fragments from PDBe, ENAMINE, and the Diamond-SGC-iNext Poised Library (DSiP) (Cox et al. 2016). Users can additionally use their own external libraries for searches. It can identify similar PDB ligands for any given ligand using pairwise PARITY comparisons (Tyzack et al. 2018) and identify compounds related to a given PDB ligand from over 35 other small-molecule database identifiers, including ChEMBL, CCDC, PubChem and DrugBank, through UniChem cross-reference service (Chambers et al. 2013), facilitating the integration and exploration of ligand information across diverse data sources.

To ensure accurate representations of complex biological ligands, such as haem, PDBe CCDUtils incorporates an enhanced data sanitization process that iteratively identifies and rectifies unusual valency issues, adjusting bond types and formal charges as necessary. To generate high-quality 2D ligand depictions, the toolkit implements a Depiction Penalty Score (DPS) to evaluate depiction quality based on bond collisions and suboptimal atom positioning. It compares both template-based and connectivity-based methods to select the best image, ensuring high-quality visual representations of ligands.

Furthermore, PDBe CCDUtils identifies CLC compounds in PDB entries, providing a more complete and precise structural representation for multi-component ligands represented using multiple CCDs with covalent links. Users can export small molecule data in various formats, including SDF, CIF, PDB, JSON, XYZ, XML, and CML. All this enriched data generated by PDBe CCDUtils is pre-computed and made available for download via FTP at https://ftp.ebi.ac.uk/pub/databases/msd/pdbechem_v2/ in the respective CCD, PRD, and CLC folders. PDBe CCDUtils is available at https://github.com/PDBeurope/ccdutils. With its comprehensive features, PDBe CCDUtils enables researchers to efficiently parse and enrich ligand data, significantly enhancing access to various ligand data in PDB. Tutorial 1.1 showcases how to read and generate enriched data for PDB’s small molecule reference dictionary. It also demonstrates how to identify and access multi-component ligand systems in a given PDB entry.

### PDBe Arpeggio: Identifying ligand interactions in PDB

Accurate identification of macromolecule-ligand interactions depends on several key factors, including appropriate protonation, complete macromolecular structure in the form of its observed biological assembly, and complete small-molecule definitions. Determining ligand protonation state is crucial for precisely defining and classifying its interactions, as protonation affects the ligand’s charge and hydrogen bonding potential. Since most PDB structures lack the resolution necessary to model hydrogen atoms directly, a protonation step must be carried out before analysing ligand interactions. Utilising the biological assembly provides a functional state of a macromolecule, capturing the full context of interactions. Additionally, considering the entire small molecule, especially in the case of multi-component ligands from the PDB, ensures that all relevant structural details are included, offering a holistic system for interaction analysis.

The PDBe interactions pipeline addresses all these factors by preparing the ligand-macromolecule system before analysis. First, the preferred biological assembly is generated using Mol* Model Server (https://molstar.org/docs/data-access-tools/model-server/). This assembly structure is then protonated using ChimeraX (Meng et al. 2023), representing the entire molecular complex rather than just the asymmetric unit for crystallographic structures. After this, complete bound molecules are inferred based on the connectivity of the non-protein chemical components (CCDs) within the assembly and each molecule is assigned a unique identifier (bmID). For each of these bound molecules, interactions are calculated utilising PDBe Arpeggio. PDBe Arpeggio is a software tool based on the Arpeggio Python library (Jubb et al. 2017), designed to calculate interatomic contacts in macromolecules, including those between ligands and various biomolecules. PDBe Arpeggio extends the original library with features specifically tailored for analysing structures in the PDB. Interatomic contacts are based on rules defined for CREDO, a protein-ligand interaction database (Schreyer and Blundell 2009), which formed the basis of the Arpeggio Python package. PDBe Arpeggio supports input files in PDBx/mmCIF format, the standard PDB archive format, and categorises interactions into four types: atom-atom, atom-plane, plane-plane, and plane-group contacts. Detailed molecular interaction information is also available through the PDBe API (https://www.ebi.ac.uk/pdbe/graph-api/pdbe_doc/#api-PDB-GetBoundMoleculeInteractions), for individual atoms in ligands and polymer residues in a PDB entry. Additionally, protonated biological assemblies are accessible via PDBe Static file (https://www.ebi.ac.uk/pdbe/static/entry/download/1cbs_bio_h.cif.gz) and are used for accurate interaction computation. PDBe Arpeggio is an open-source python package available at https://github.com/PDBeurope/arpeggio. The tutorial 1.2 presents how the PDBe interactions pipeline prepares the structure and uses PDBe Arpeggio to calculate macromolecule-ligand interactions.

### PDBe RelLig: Identifying the functional role of ligands in PDB

Identifying ligand roles in protein structures is key to understanding macromolecular functions, enzyme mechanisms, and drug interactions (Westbrook and Burley 2019; Burley et al. 2019; Credille et al. 2019; Moreira et al. 2019; Richard 2022; Vetting et al. 2015). While cofactors and substrates reveal protein activity, others, like buffer components or cryoprotectants, may be introduced during experimental procedures (Peat, Christopher, and Newman 2005; Caffrey and Cherezov 2009; McPherson and Cudney 2014; Garman 2003; Pflugrath 2015; Jang et al. 2022). Distinguishing functional ligands from non-functional ones is crucial for accurate biological interpretation. The PDBe RelLig (Relevant Ligands) pipeline addresses this by identifying biologically relevant ligands and categorising them based on their functional roles in the entries where they are bound—such as cofactor-like, reactant-like (similar to substrate or product), or drug-like—placing them within their biological context (Figure 3).

**Figure 3.**
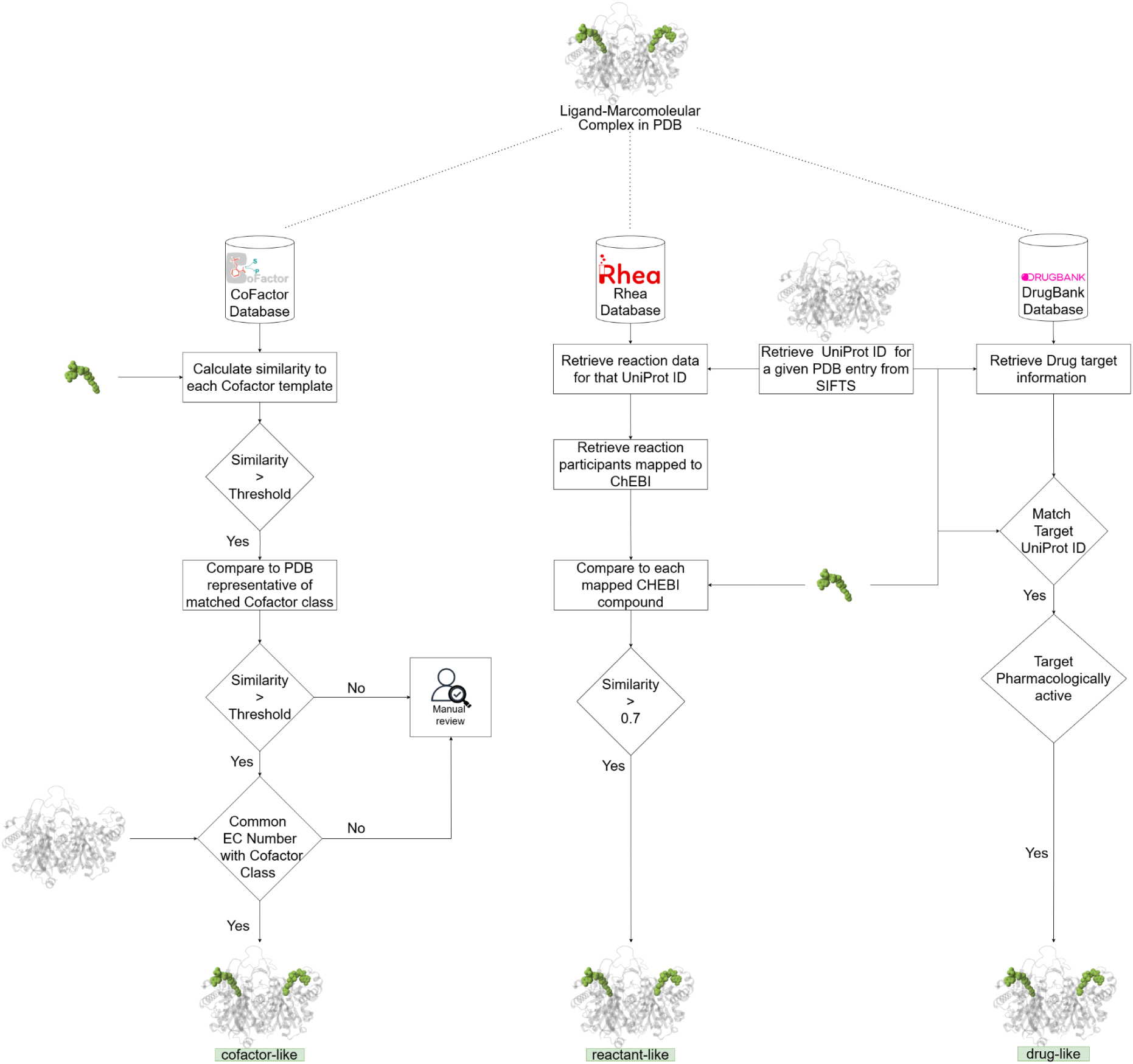
Schematic overview of the PDBe RelLig Pipeline. This schematic outlines the identification of relevant ligands in the PDB and their categorisation based on functional roles within protein structures. Ligands are classified as cofactor-like, reactant-like, or drug-like.

### Identifying cofactors-like ligands in PDB

The PDBe RelLig pipeline identifies cofactor-like ligands in the PDB using a semi-automated annotation process based on the 2D structural similarity of ligands to cofactor classes in the CoFactor database (Fischer, Holliday, and Thornton 2010). The CoFactor database contains 27 classes of manually curated organic enzyme cofactors and details of associated enzymes including EC numbers. For each cofactor class, we identified a small molecule from the PDB closely matching the structure of the template molecule based on PARITY similarity and assigned it as a representative molecule (Mukhopadhyay et al. 2019). Additionally, we defined a minimum threshold for each cofactor and extended the list of enzyme EC numbers associated with the cofactor classes using the information from BRENDA (Chang et al. 2021). Using this information, for each newly released small molecule in the PDB, similarity is calculated against the template molecules of each cofactor class. If the similarity meets the minimum threshold of any cofactor class, the ligands are further evaluated against the representative molecule for the matched cofactor class. When the similarity score remains above the threshold and the ligand is present in a PDB entry with an approved EC number corresponding to the cofactor class, the ligand is classified as cofactor-like (Mukhopadhyay et al. 2019). Ligands that do not meet these criteria are flagged for manual annotation, ensuring accurate identification of biologically relevant ligands.

### Identifying reactant-like ligands in PDB

The PDBe RelLig pipeline identifies reactant-like ligands in the PDB by mapping to the Rhea database (Bansal et al. 2022), an expert-curated resource that uses the ChEBI ontology (Hastings et al. 2016) to describe reaction participants and their structures. For each reaction in Rhea, all associated PDB structures are mapped to the UniProt accession of the catalysing protein. Using the PARITY method, the bound ligands in these PDB structures are compared to ChEBI compounds involved in the reaction. Ligands that achieve a minimum similarity score of 0.7 are annotated as reactant-like, ensuring accurate identification of biologically relevant reactants (substrate or product of the reaction).

### Identifying drug-like ligands in PDB

The PDBe RelLig pipeline identifies drug-like ligands in the PDB by mapping them to the DrugBank database (Knox et al. 2024). Ligands bound to PDB structures of pharmacologically active targets listed in DrugBank are classified as drug-like, enabling the accurate annotation of ligands with potential therapeutic relevance. This approach ensures precise identification of biologically and pharmacologically significant drug-like ligands in the PDB.

The PDBe RelLig pipeline is open-source and available at https://github.com/PDBeurope/rellig. Tutorial 1.3 demonstrates how the PDBe RelLig pipeline utilises the PARITY method to identify and classify ligands in PDB structures as reactant-like, cofactor-like, or drug-like.

### PDBe web pages for ligand visualisation and analysis

For any given structure in the PDB, the “Ligands and Environments” section provides a detailed overview of bound ligands, cofactors, and modified residues, as seen in PDB ID 4p68, which includes one cofactor, three ligands, and one modified residue. By clicking on the ligand image or ligand identifier (e.g., CCD ID: NAD), an interactive visualisation is launched, showcasing PDBe Arpeggio calculated residue-level interactions in 2D using the LigEnv web component and atomic-level interactions in 3D between ligands and their binding sites using Mol* (Figure 4). This dual-view visualisation is intercommunicative—if a user clicks on a residue in the 2D schematic, it is highlighted in the 3D viewer and vice-versa. This integration facilitates easy analysis of ligand-protein interactions by visualising spatial and molecular relationships intuitively. The LigEnv web component ensures that ligands are presented in a consistent orientation and layout across different PDB entries, facilitating comparison of the same ligands across various binding sites. Residues forming the ligand binding site are colour-coded according to their properties, improving clarity and enhancing the user’s ability to interpret the interactions. Users can export the underlying interaction data in JSON format for computational analysis or export visual representations in SVG format for documentation. The LigEnv component is an open-source web component available at https://github.com/PDBeurope/ligand-env/.

**Figure 4.**
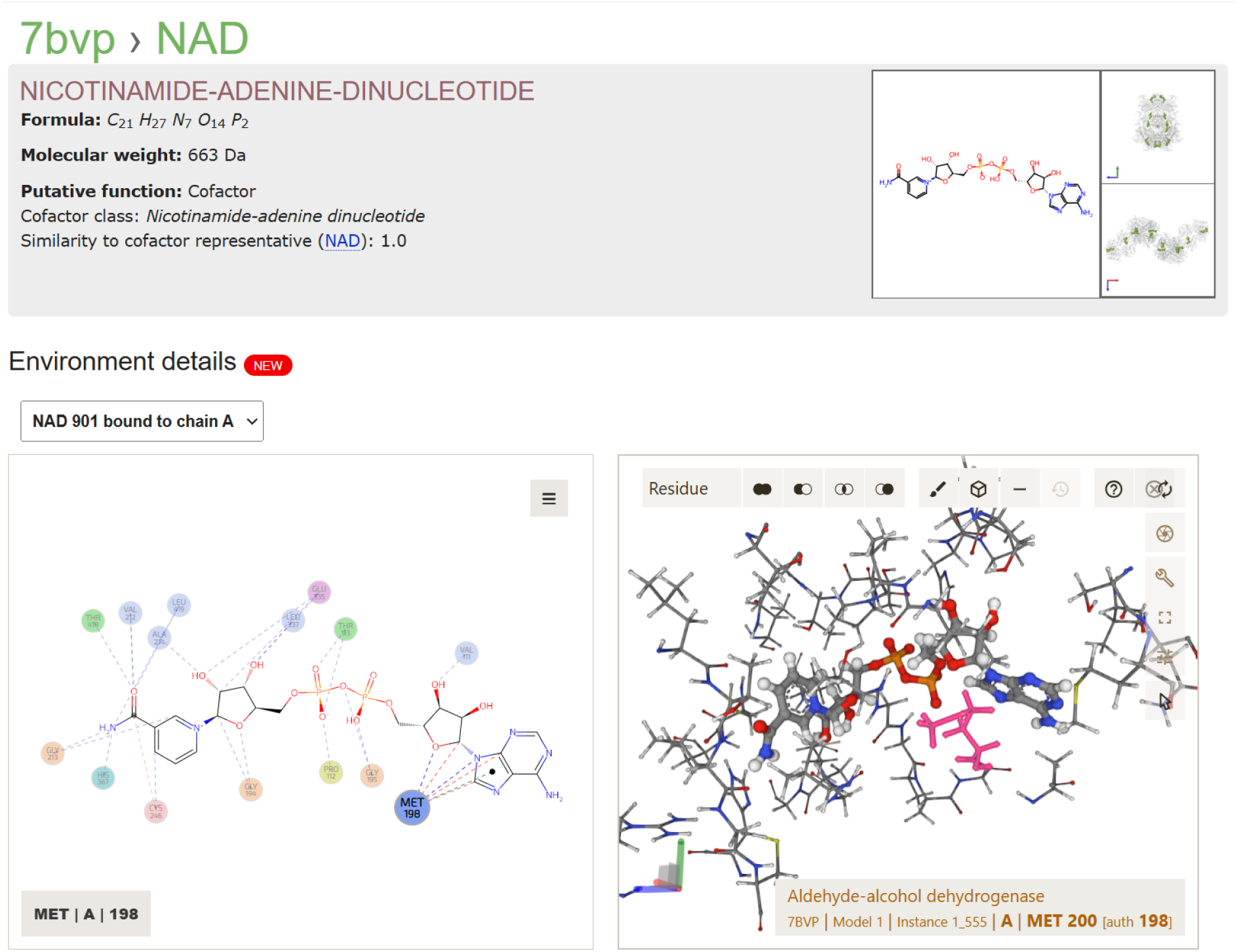
Cofactor Binding Site of Aldehyde-alcohol Dehydrogenase (PDB ID: 7BVP). The bound ligand NAD is annotated as cofactor and required for the reduction of the carbonyl group. The ligand binding residue Methionine 198 in LigEnV web component is automatically highlighted in the Mol* viewer allowing for easy identification of key binding residues in the 3D structure.

Additionally, the wwPDB validation reports (Gore et al. 2017; Feng et al. 2021), accessible through PDBe entry pages, provide essential information on the quality of ligands (Figure 5). Using the program Mogul (Bruno et al. 2004), these reports evaluate ligand geometry against the small molecule structures in the Cambridge Structural Database (CSD) (Groom et al. 2016; Ferrence et al. 2023). Z-scores are calculated to quantify deviations in bond lengths, angles, torsion angles, and ring geometry, with values above 2.0 flagged as outliers. The RMSZ score, which summarises the overall ligand geometry, should ideally range between 0 and 1. For torsion angles, deviations are flagged if their local density measure is below 5%, while rings are flagged if their torsion angle RMSD exceeds 60°. Chirality and stereochemistry are also assessed for errors. Beyond geometric validation, ligand fit to the electron density is assessed using metrics like the RSR (Real-Space R-value) and RSCC (Real-Space Correlation Coefficient). Ligands with RSCC values below 0.8 or RSR values above 0.4 are highlighted for further scrutiny (Smart et al. 2018).

**Figure 5.**
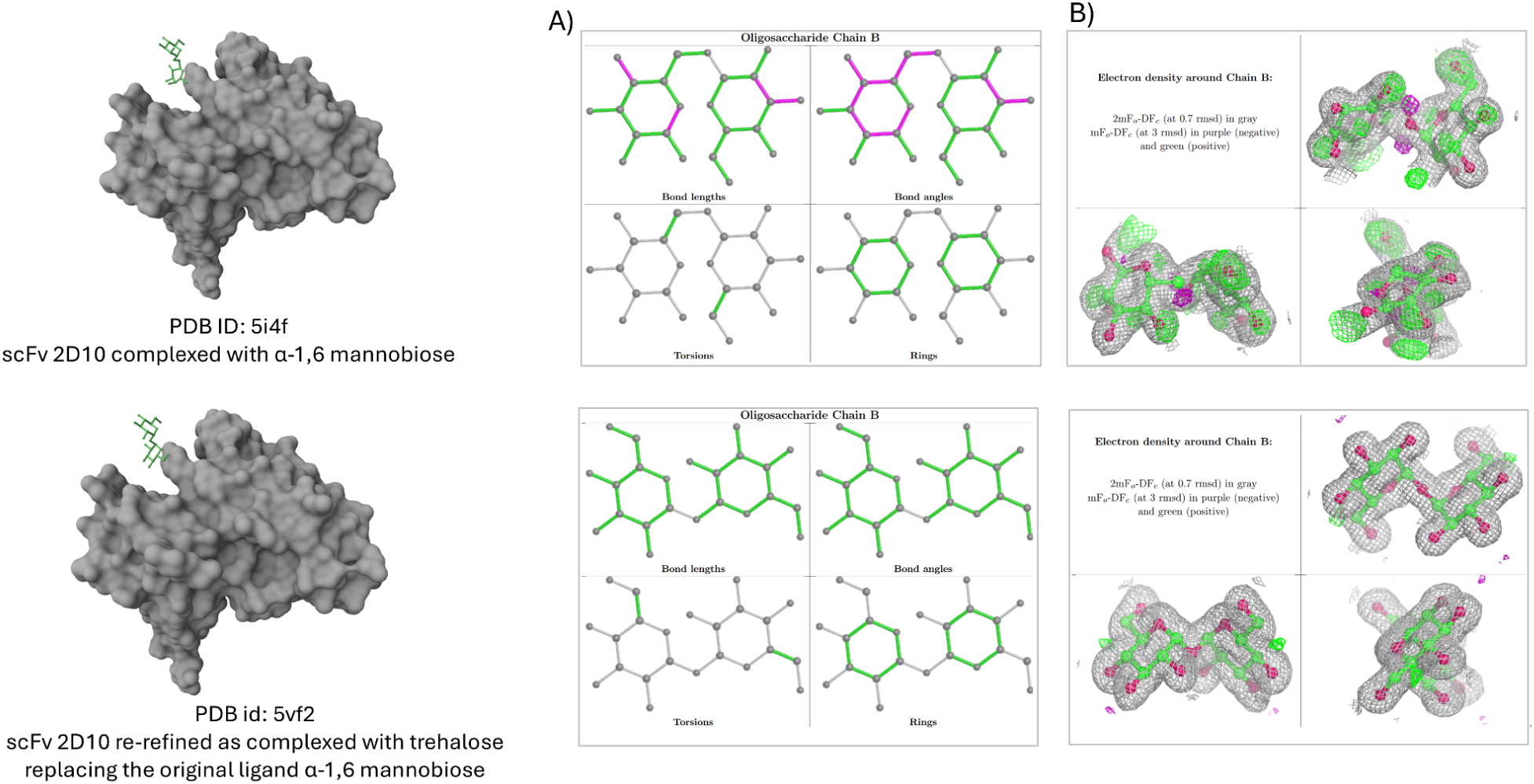
Ligand validation assessment of scFv 2D10 Structures. Comparison of the original structure bound to α-1,6-mannobiose (PDB ID: 5I4F) and the re-refined structure bound to trehalose (PDB ID: 5VF2) at the same binding site. A) 2D image of ligand geometric quality. Commonly observed values are shown in green, unusual values in magenta, and features with insufficient data in grey. B) The 3D view for the ligand atomic model fit the experimental electron density map (shown in grey) with positive and negative difference density maps shown in green and magenta, respectively. The re-refined structure exhibits no geometric outliers and a significantly better fit, indicating improved model quality and confirming the misidentification of the principal ligand in the original structure.

These validation tools help identify potential issues with ligand modelling, allowing users to compare ligand quality across models and select those with better fit and geometry for more accurate structural representation. Tutorial 2.1 demonstrates how to examine PDB-ligand interactions on PDBe entry pages and assess the quality of ligands using wwPDB validation reports.

### PDBe-KB: Enabling ligand data analysis in its biological context

The PDBe-Knowledge Base (PDBe-KB) , established in 2018 and maintained by the PDBe team at EMBL-EBI, is an open and collaborative data resource dedicated to placing macromolecular structure data within their biological context (PDBe-KB consortium 2022). PDBe-KB consortium partner resources contribute biochemical, biophysical, and functional annotations based on analysis of the PDB structures. As of 2024, PDBe-KB has integrated contributions from 34 consortium partners, including numerous cheminformatics experts who contribute data on small molecules and their macromolecular binding sites, with collaborators such as canSAR(di Micco et al. 2023), 3DLigandSite (McGreig et al. 2022), P2Rank (Krivák and Hoksza 2018), and many others, as detailed at pdbe-kb.org/partners. These annotations, encompassing curated and predicted data, are crucial for understanding the roles and mechanisms of small molecules interacting with proteins and nucleic acids. Examples of these annotations include catalytic sites, biologically relevant metal ions, druggable pockets, and predicted ligand binding sites. Table 2 provides more details on various ligand resources available in PDBe-KB.

**Table 2:**
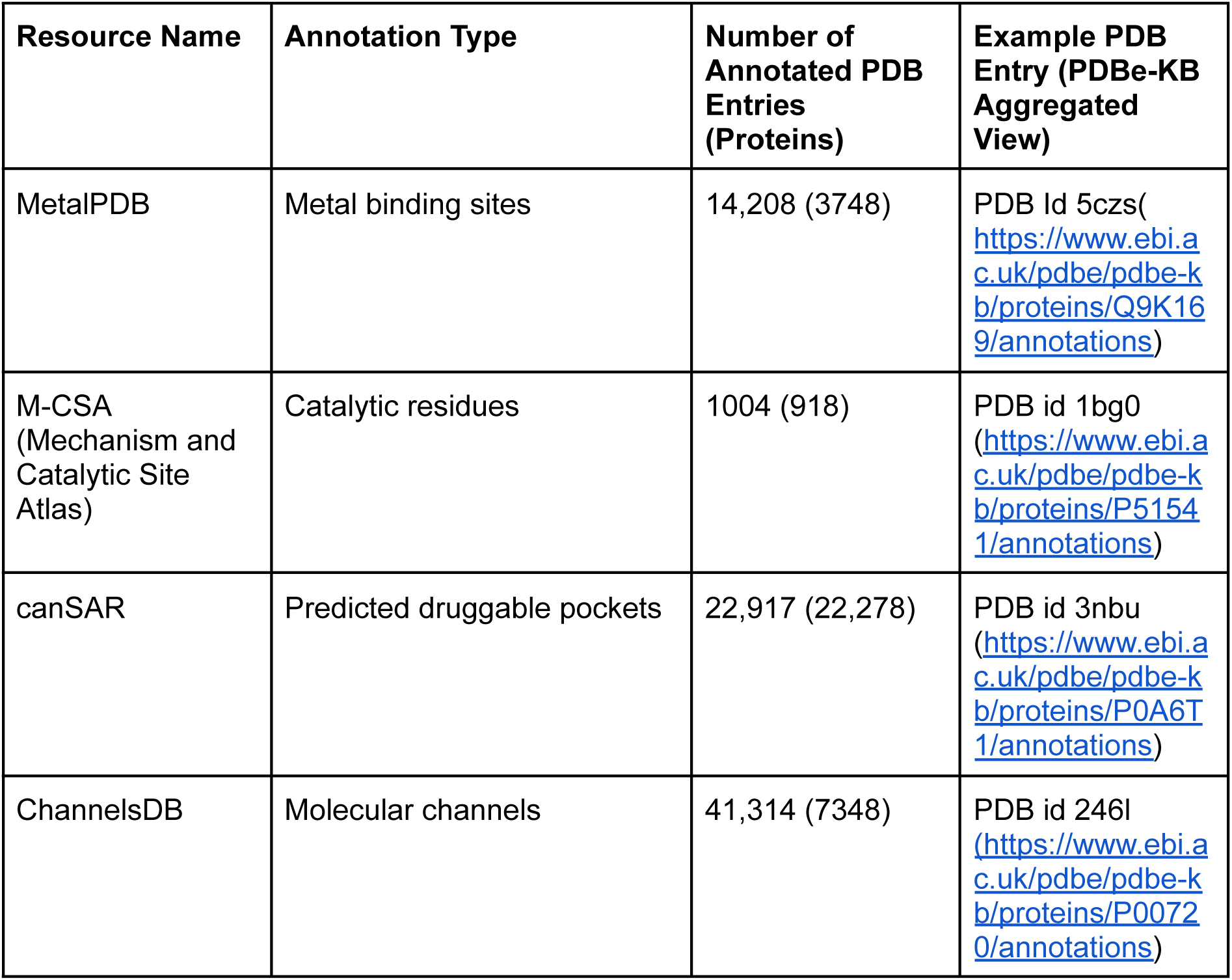
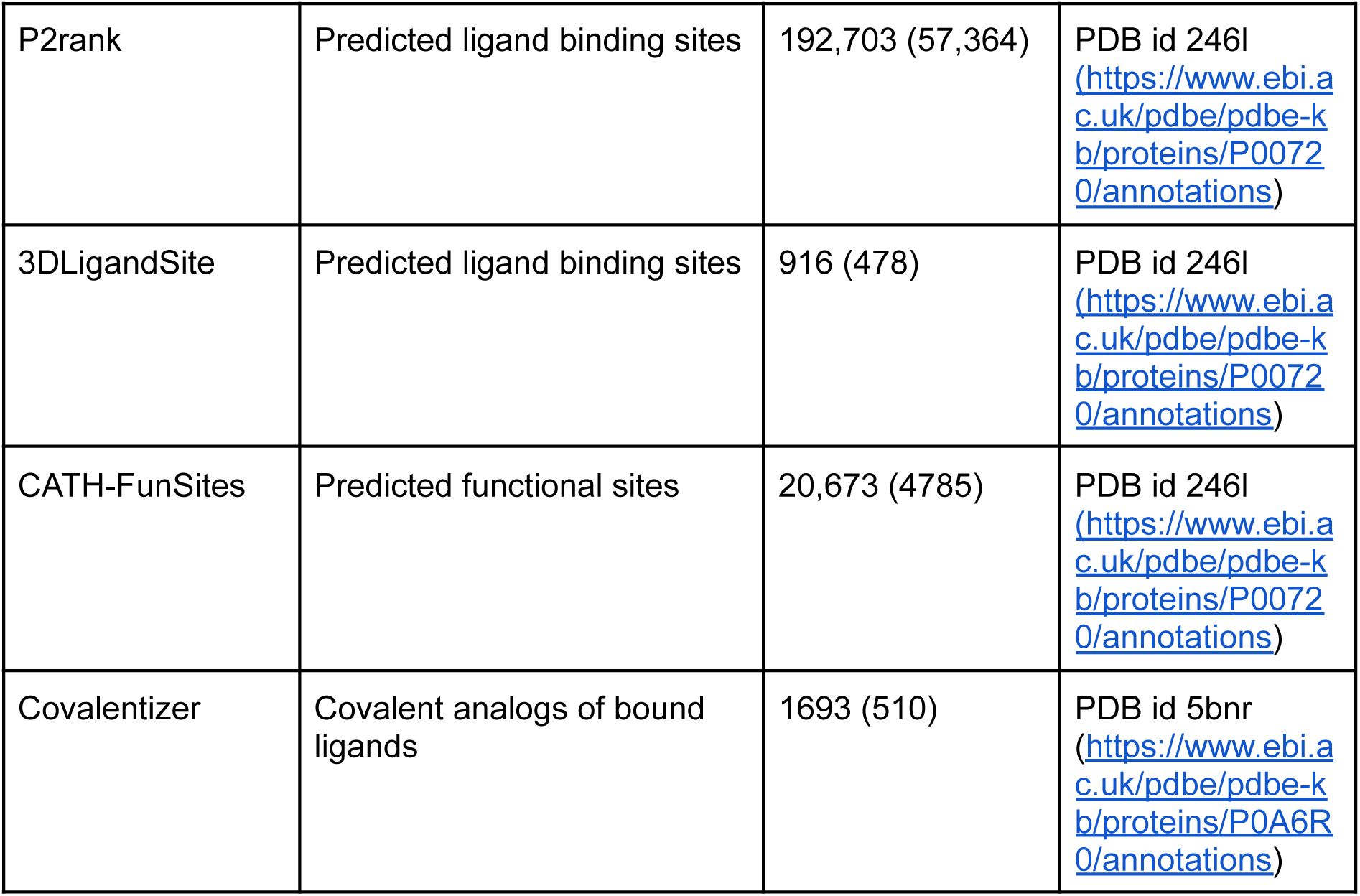
Small-molecule-related resources and annotations in PDBe-KB.

PDBe-KB aims to consolidate the wealth of information and enable efficient use of structure data by aggregating all the data in a knowledge graph for a given biological entity (i.e. complexes, proteins, domains, ligands). The aggregated protein view, the first PDBe-KB webpages, was launched in March 2019, and is being actively developed with new annotations and features.

### Ligand annotations on PDBe-KB protein pages

The PDBe-KB aggregated views of proteins integrate data for a given protein (based on common UniProt accession) from multiple PDB entries to present a comprehensive web-based visualisation of all the relevant structural data. These web pages feature a sub-page showcasing an integrated view, “Ligands and Environments”, of all small molecules from all relevant entries from the PDB archive that interact with the given protein, along with annotations regarding their functional roles, such as drugs, cofactors or reactants identified using the PDBe RelLig pipeline (Figure 6A). The ligands are organised based on their chemical scaffolds, enhancing visual comparison and accessibility. Each entry in the gallery displays the chemical structure, its CCD ID, recommended name, and additional information. Users can explore detailed 3D representations of ligand interactions through the Mol* 3D viewer by clicking on the ligand image and downloading relevant PDB entries in CSV or JSON formats using the download button.

**Figure 6.**
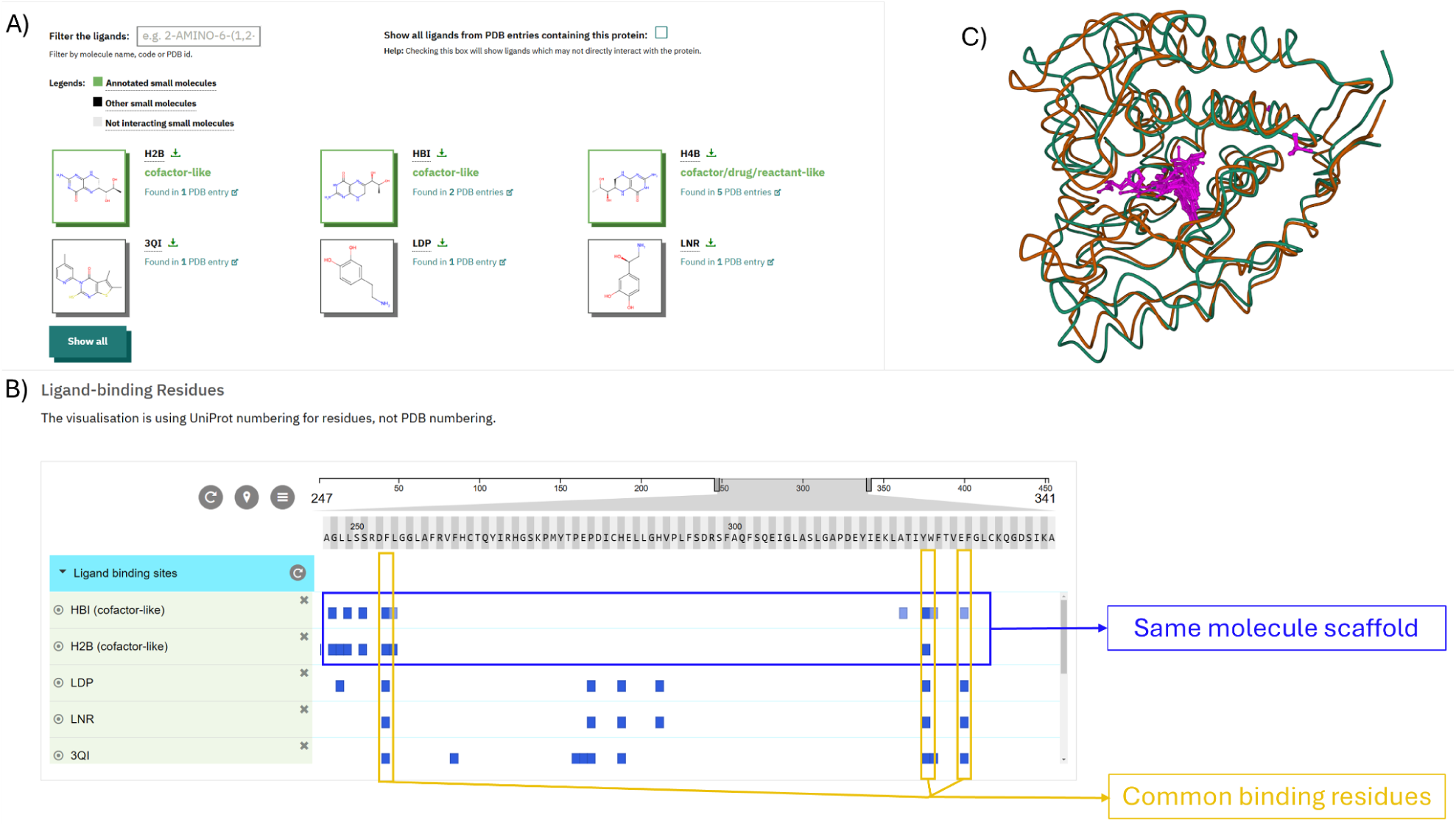
Ligand data for Phenylalanine-4-hydroxylase (P00439) on PDBe-KB Protein Page. A) Ligand gallery displaying all ligands bound to Phenylalanine-4-hydroxylase, with cofactors highlighted in green. B) PDBe ProtVista viewer displaying ligand-binding residues, facilitating easy identification of common binding residues and those specific to cofactors H2B, HBI, and H4B, which share a common molecular scaffold. C) 3D superposition of ligands, highlighting the most prevalent binding pockets.

The “Ligands-binding Residues” section provides a comprehensive view of ligand binding residues from all relevant PDB entries containing the given protein, using a 2D sequence viewer, PDBe ProtVista (Figure 6B). This viewer uses darker shades of blue to indicate frequently interacting residues, enabling researchers to gain insights into key binding residues and sites. A collapsed view provides a summary that can help in identifying binding sites that bind a diverse set of small molecules.

Furthermore, users can access a superposed view of small molecules from various PDB entries for specific protein residue ranges(“segment”), highlighting all binding pockets. By clicking the “3D view of superposed ligands” button, users can visualise representative conformations for a given protein (PDBe-KB consortium 2022; Ellaway et al. 2024) alongside all ligands associated with that segment (Figure 6C), which is particularly beneficial for analysing proteins identified in fragment screening experiments. Collectively, these features enhance the understanding of ligand-protein interactions and support more accurate structural representation and analysis. Tutorial 2.2 illustrates how to identify key binding residues on PDBe-KB aggregated views of proteins and locate important ligand binding sites using a 3D view of superimposed ligands.

### PDBe-KB ligand pages

In the next phase of PDBe-KB development, an aggregated view for individual ligands has been created to consolidate data on small molecules from the Protein Data Bank (PDB) and provide enhanced biological context. By leveraging the integrated PDBe-KB knowledge graph, this view simplifies the traditionally labour-intensive process of linking and analysing macromolecular interactions with small molecules. These ligand-centric pages, keyed on PDB ligand identifiers such as CCD, PRD and CLC IDs, present a comprehensive overview of each ligand’s chemical, structural, interaction and functional information in a user-friendly and intuitive manner.

The essential ligand features, including molecular name, synonyms, molecular formula, and chemical descriptors such as IUPAC InChI, InChIKey, and SMILES, are displayed in the “Description” tab. Structural representations are provided in 2D and 3D formats, with distinct views highlighting the overall ligand structure, individual atoms, the Murcko scaffold, and specific ligand fragments. Physicochemical properties, categorised into molecular, ring, conformational, surface, functional group, and stereochemical properties, are also shown. Additionally, if the CCD is part of a larger BIRD (PRD) or CLC entry, this information is also shown here. All data in this tab is generated using PDBe CCDUtils.

The “Structures” tab provides access to all PDB structures, where ligand-protein interactions for a given ligand are observed, in a table format. The table includes key details such as the protein name, UniProt accession (with link to PDBe-KB aggregated view for that protein), total number of PDB structures, source organism, EC number, and the functional role of the ligand (e.g., drug-like, cofactor-like or reactant-like) as determined by the PDBe RelLig pipeline. Since the same protein may have multiple PDB structures resolved under different experimental conditions or with mutations, the data is grouped by default in the “Proteins” view, showing the total number of PDB structures. Users can switch to the “Structures” view to see all the individual PDB entries and easily find relevant protein-ligand complexes.

The “Interaction statistics” tab aggregates atom-level interaction data for all instances of the ligand bound to proteins in the PDB. A heatmap displays the relative interaction frequency for each ligand atom with respect to total number of interactions by the ligand, with darker colours indicating more frequent interactions (Figure 7). This data is also mapped onto a 2D ligand image, providing an overview of interaction hotspots. The heatmap and ligand image are interactive, allowing users to highlight corresponding atoms across both views simultaneously. Additionally, the heatmap shows the likelihood of a given ligand atom to interact with specific amino acids, where interaction frequency can be sorted by either amino acid type or interaction frequency, offering insights into whether a ligand preferentially interacts with certain types of residues and allowing for quick identification of the most frequently interacting amino acids. Interaction data can also be filtered by interaction types, such as hydrogen bonds, Van der Waals, polar, or hydrophobic interactions.

**Figure 7.**
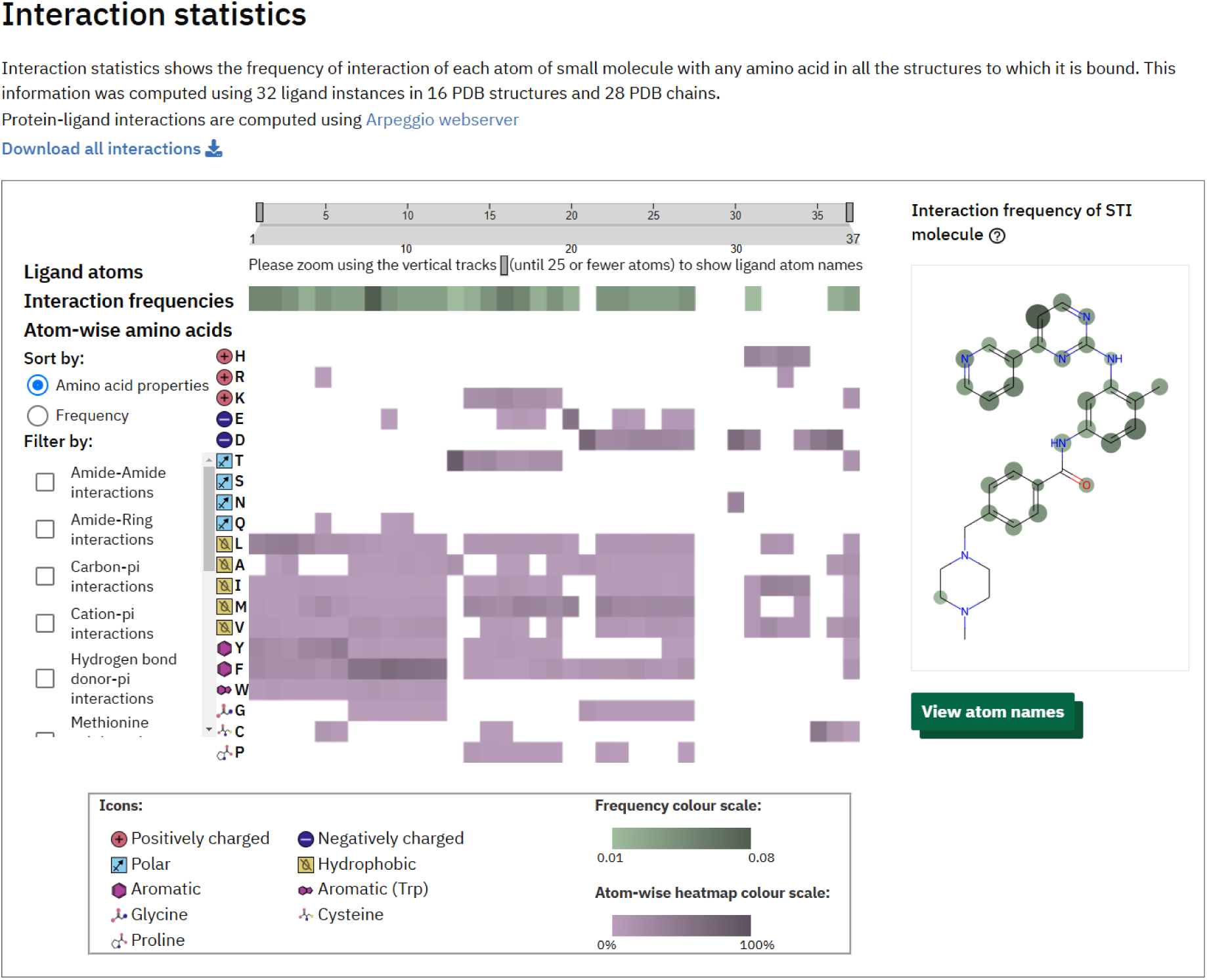
Aggregated interaction patterns for Imatinib (CCD ID: STI) on PDBe-KB ligand pages. Imatinib primarily interacts with hydrophobic residues, along with a few aromatic residues as illustrated on the heatmap. Key interaction hotspots for Imatinib are highlighted on its 2D image, showing that interactions primarily occur through its phenyl, pyridine, and pyrimidine rings.

The “Related ligands” tab provides access to structurally similar ligands, identified using the PDBe RelLig pipeline. The similar ligands are organised into three sub-sections based on the type of similarity: ligands with the same scaffold, those with ≥60% similarity, and any stereoisomers. Each sub-section features an image gallery displaying the ligand structures, with a 3D view of the ligand upon clicking an image. Additionally, details such as the ligand name, CCD ID, percentage similarity, and the number of PDB structures in which the similar ligands are found are provided, along with a link to the PDBe search page. This search page lists all relevant PDB structures and includes various filtering options on the left menu, such as experimental method, source organism, sequence, structural classification, etc.

Finally, the “Ligand-specific databases” offers cross-references to other small-molecule data resources generated using UniChem integration in PDBe CCDUtils. A data download functionality is integrated within each section for easy access to data for further analysis. By integrating chemical, structural, interaction, and functional data, the PDBe-KB Ligand Pages provide an aggregated view on small molecules, enhancing the accessibility and biological relevance of ligand data within the PDB. Tutorial 2.3 showcases how to answer scientific questions using PDBe-KB ligand pages.

### PDBe-KB programmatic data access

All the data displayed on PDBe and PDBe-KB web pages are powered by REST API endpoints, enabling users to access this information programmatically (Nair et al. 2021). These API endpoints are organised into categories such as compounds, proteins (UniProt), residues, PDB entries, and validation for efficient navigation. The compound-related endpoints provide information on all PDB entries containing specific ligands, details about their atoms and bonds, similar ligands, ligand substructures, and functional annotations. Additionally, they offer summary data, including descriptors such as InChI, InChIKey, SMILES, and physicochemical properties. A complete list of API endpoints is available at https://www.ebi.ac.uk/pdbe/graph-api/pdbe_doc. These API endpoints query the PDBe graph database to retrieve the necessary information. The PDBe graph database is freely available for download at https://www.ebi.ac.uk/pdbe/pdbe-kb/graph, enabling users to conduct advanced queries and custom analyses. Tutorial 3 illustrates how to programmatically access ligand data in PDBe and PDBe-KB through various API endpoints.

### Summary Files for Comprehensive Analysis

PDBe offers summary files to facilitate extensive analyses of ligand interactions in the PDB archive. Two essential files are Interacting_chains_with_ligand_functions.tsv and pdb_bound_molecules.tsv are available at https://ftp.ebi.ac.uk/pub/databases/msd/pdbechem_v2/additional_data/pdb_ligand_interactions/. The first file summarises ligand interactions, detailing interacting macromolecule chains, their UniProt accessions, functional annotations, and identifiers such as InChIKey, bmID, and LigandType. The second file contains information on CLC molecules and their composition.

The ChEMBL and PDBe teams have mapped over 17,000 PDB-ligand complexes to approximately 39,000 bioactivity records, encompassing various experimental data like binding affinities and functional assays. These results are available at https://ftp.ebi.ac.uk/pub/databases/msd/pdbechem_v2/additional_data/bioactivity_reports/, including a comprehensive report and a simplified version with aggregated values for each target-ligand complex and direct links for further exploration of ChEMBL data.

## Discussion

Identifying and accurately representing ligands is crucial for understanding protein-ligand interactions, which have wide-ranging applications in understanding protein function, target validation, drug development, and repurposing. With the introduction of CLCs, PDBe enables chemically complete and precise representations of multi-component ligands. The accurate identification and data standardisation of small molecules via PDBe CCDUtils has enabled the mapping of many previously overlooked small molecules from the PDB to other databases. This process has significantly enhanced the completeness and accuracy of ligand definitions and facilitated the integration of ligand data across these resources and as a result facilitates for example the identification relevant ligand-target complexes. The PDBe RelLig pipeline provides annotations that clarify the functional roles of ligands in their targets, helping researchers distinguish biologically significant molecules from experimental artefacts and streamlining the analysis process.

Moreover, the integrated information on PDBe ligand pages enables users to view small molecule data within a comprehensive structural context. This holistic view encompasses ligand descriptions, physicochemical properties, ligand-protein complexes, and the functional roles of ligands within those complexes. Additionally, it provides overall binding interaction statistics, related ligands, and cross-links to other small molecule databases. This wealth of information aids researchers in identifying critical patterns in ligand interactions, understanding these molecules’ biological significance, and exploring potential applications in various fields, such as drug discovery, enzyme design, and biomolecular research.

## Conclusions

The PDB contains a wealth of information on small molecules and their interactions with macromolecules, but specialised tools are essential to fully leverage this data. PDBe has developed several open-access tools—PDBe CCDUtils, PDBe Arpeggio, and PDBe RelLig—that enrich ligand data, enable detailed interaction analysis, and classify ligands based on their biological roles. The PDBe-KB ligand pages also provide an integrated platform for accessing and visualising this enriched data. Together, these resources empower researchers to efficiently explore the properties of small molecules, analyse their interactions with proteins, and distinguish between biologically relevant ligands and experimental artefacts. By integrating these tools and resources into the PDBe ecosystem, we provide a comprehensive platform for analysing and visualising small molecules within the PDB, greatly enhancing understanding of their biological context and functional significance to facilitate basic and translational research.

## Acknowledgements

The authors would like to express their gratitude to the UKRI-Biotechnology and Biological Sciences Research Council for funding provided under the BioChemGraph project (BB/T01959X/1) and to the European Molecular Biology Laboratory-European Bioinformatics Institute for their support. Special thanks are extended to collaborators from ChEMBL: Melissa F. Adasme, James Blackshaw, and Andrew Leach, as well as from CCDC: David Lowe and Ian Bruno.

## Author’s Contribution

**Preeti Choudhary**: Project administration (lead); conceptualisation (equal); supervision (lead); software (equal); formal analysis (equal); visualisation (lead); writing – original draft (lead). **Ibrahim Roshan Kunnakkattu**: software (lead); methodology (lead); formal analysis (equal); visualisation (equal); writing – review and editing (equal). **Sreenath S Nair**: software (equal); supervision (equal). **Dare Kayode Lawal**: software (equal); visualisation (equal). **Ivanna Pidruchna**: visualisation (equal). **Marcelo Querino Lima Afonso**: software (supporting), visualisation (supporting). **Jennifer Fleming:** supervision (equal); conceptualisation (equal); writing – review and editing (equal). **Sameer Velankar**: conceptualisation (equal); funding acquisition (lead); investigation (equal); resources (lead); supervision (equal); writing – review and editing (equal).

## Notes

https://github.com/PDBeurope/ccdutils

https://github.com/PDBeurope/arpeggio

https://github.com/PDBeurope/rellig

https://wwwdev.ebi.ac.uk/pdbe-srv/pdbechem/chemicalCompound/show/STI

https://github.com/PDBeurope/pdbe-notebooks/blob/main/pdbe_ligands_tutorials/PDBe_ligand_tools.ipynb

https://github.com/PDBeurope/pdbe-notebooks/blob/main/pdbe_ligands_tutorials/PDBe_PDBe-KB_ligands_webpages_tutorials.pdf

https://github.com/PDBeurope/pdbe-notebooks/blob/main/pdbe_ligands_tutorials/PDBe_KB_API_tutorials.ipynb

### Competing Interest Statement

The authors have declared no competing interest.

